# The Diversity of NLRs in *Brassica rapa* Pan-genome

**DOI:** 10.1101/2023.08.29.555307

**Authors:** Shuhao Liu, Yimeng Zhu, Boyang Ke, Jun Wang, Xianjia Peng, Shaoke Zhang, Ruoxuan Zhao, Haizheng Wang, Jing Lou, Jiaruo Li, Yunhong Zheng, Tao Huang, Siyao Wang, Zebin Lang, Leye Wang, Xizhe Sun, Chunyang Wang

**Affiliations:** College of Life Sciences, Hebei Agricultural University, Baoding, Hebei, 071001, China; State Key Laboratory of North China Crop Improvement and Regulation, Hebei Agricultural University, Baoding, Hebei, 071001, China; Ministry of Education of China-Hebei Province Joint Innovation Center for Efficient Green Vegetable Industry, Baoding, Hebei, 071001, China; Hebei Bioinformatic Utilization and Technological Innovation Center for Agricultural Microbes, Baoding, Hebei, 071001, China; School of Life Sciences, Hebei University, Baoding, Hebei, 071002, China

## Abstract

The species *Brassica rapa* is mainly cultivated as a vegetable crop with high economic value. But infectious diseases, such as soft rot disease and clubroot disease can cause severe yield losses. Introducing resistance genes through breeding is an efficient and sustainable way to reduce the susceptibility and improve the yield of crops. Nucleotide-Binding Leucine-Rich Repeat (NLR) genes are the main types of resistance genes. They can recognize pathogen effectors and initiate downstream immune response. However, the intraspecific diversity of *NLR* genes in *Brassica rapa* has remained unknown. Based on domain similarity search and machine learning pipeline, we identified 2188 NLRs in 17 accessions of *Brassica rapa* genomes, which constitute the species-wide pan-NLRome in *Brassica rapa*. The diversity of the four types of *NLR* genes are significantly different in the aspects of chromosome location, number stability, integrated domains (IDs), evolutionary trajectory, and positive selection sites. Phylogenetic analyses show TNL-type NLRs whose N terminus contain Toll/interleukin-1 receptor (TIR) domain experienced accession-specific expansion. Moreover, the expanded TNL-type NLRs carry more positively selected amino acid residues which are mainly located on the protein surface based on their 3D structures. These evidences might imply their important biological function in perceiving pathogen effectors. Taken together, our study provides better insights into the diversity and variation of *NLR* genes in *Brassica rapa* pangemone, which can facilitate the biofunction assigning, cloning, and subsequent application in breeding programs.

## Introduction

*Brassica rapa* (AA, 2n=20) is a species of *Brassica* genus belonging to Brassicaceae family. It is widely cultivated throughout worldwide and is rich in nutrients (Cai et al., 2021). As the main sources of vegetables, *Brassica rapa* has been domesticated into different subspecies and varieties, such as oil seed, Chinese cabbage, and so on. For example, Chinese cabbage (*Brassica rapa* ssp. *pekinensis*) is the most important vegetable in Asia (Sun et al., 2022). However, many bacteria, fungi, and oomycete pathogens can attack *Brassica rapa* crops. They can cause serious diseases, such as soft rot (*Pectobacterium carotovorum* ssp. *carotovorum*), blackleg (*Leptosphaeria maculans*), clubroot (*Plasmodiophora brasicae*), downy mildew (*Hyaloperonospora parasitica*) (Liu et al., 2019; Tirnaz et al., 2020). To the *Brassica* species, these diseases can cause 15-20% yield loss on average, while in some years, the losses can reach 90% (Chen et al., 2021; Zhang et al., 2020).

The Nucleotide-Binding Rich Leucine Repeat (NLR) genes are the largest family of plant disease resistance genes (Van Ghelder et al., 2019). In plants, as the immune receptors within cells, NLR proteins are able to recognize pathogenic proteins and their effectors on the host (Gao et al., 2018; Wang et al., 2020; Wang et al., 2023a). Plant genomes encode multiple disease resistance proteins which are essential to trigger and increase the immune response for pathogens (Tang et al., 2017). Genetic recombination might be the genetic basis for the diversity of *NLR* genes (Lal et al., 2020). Though with the high copy number, *NLR* genes are highly conserved in structure. In a protein encoded by *NLR* gene, the nucleotide-binding site (NBS) domain is located in the middle and the leucine-rich repeat (LRR) domain is located at the C-terminus (Xiao et al., 2001; Shao et al., 2014). There are four types of NLR proteins according to the N-terminal domains: the TIR-NBS-LRR type NLR proteins (TNLs) with a Toll/IL-1 receptor (TIR) domain, the CC-NBS-LRR type NLR proteins (CNLs) with a coiled-coil (CC) domain, the RPW8-NBS-LRR type NLR proteins (RNLs) with a RESISTANCE TO POWDERY MILDEW8 (RPW8) domain, and NBS-LRR type NLR proteins (NLs) without the above mentioned three domains (Shao et al., 2019; Tirnaz et al., 2020; Van de Weyer et al., 2019). Different type NLR proteins might play different roles in the immune response for pathogens. Previous studies showed that CNL-type and RNL-type NLRs can form oligomeric resistosomes and then conduct Ca^2+^ to trigger immune responses (Bi et al., 2021; Jacob et al., 2021). Such as CNL-type protein ZAR1 of *Arabidopsis* can form resistosome and plays function as a calciumpermeable channel for effector-triggered immunity (ETI) signaling (Bi et al., 2021). TNL-type protein RPP1 of *Arabidopsis* and Roq1 from tobacco also can form resistosome and function as nicotinamide adenine dinucleotide hydrolase (NADase) enzymes confered by TIR domain (Horsefield et al., 2019; Wan et al., 2019). In addition, some NLR proteins carry integrated domains (IDs) which can assist them in perceiving effectors indirectly or directly (Kourelis and Van der Hoorn 2018). The NLR proteins with IDs (NLR-IDs) might be the result of the effector target integration with the canonical NLR structure (Marchal et al., 2022). All the studied IDs might have the possibility to reflect the original targets of the pathogens so far, which indicates that they can be used as tool kits to identify the original effector targets in the host cell (Michalopoulou et al., 2022).

With the development of biochemical techniques and advanced genetic manipulation tools, several *NLR* genes have been cloned and functionally characterized in *Brassica rapa* crops, such as *Bra_cRa/cRb* and *Bra_Crr1a* (Cantila et al., 2022). Moreover, these two genes all can encode TIR-NBS-LRR proteins (*BraA08g039211E* and *BraA08g039212E*) and are identified as the most likely clubroot-resistant candidates (Wang et al., 2022). However, the cultivars containing clubroot resistant genes have been attacked by *PbXm*, a newly evolved *Plasmodiophora brassicae* isolates (Wang et al., 2023b). In order to cope with rapidly evolved pathogens, more *NLR* genes in *Brassica rapa* genomes are needed to be systematically mined and applied in breeding programs to produce improved varieties. Here, we carried out domain similarity search and machine learning combined pipeline to mine *NLR* genes in 17 different accessions of *Brassica rapa* (Cai et al., 2021; Supplemental Table S1). The identified *NLR* genes have great application values. Extensive excavation of *NLR* genes is an important way to select and breed disease-resistant varieties, because disease-resistant breeding is the most effective method of disease control. Cultivating disease-resistant varieties has reduced the use of chemical drugs to a certain extent, alleviating environmental pollution and food health problems. Our study establishes foundations for the characterization and functional study of *NLR* genes in *Brassica rapa* crops.

## Results

### Unequal distribution of *NLR* genes

In the 17 *Brassica rapa* accessions, we found a total of 2188 NLRs, including 768 NLs, 702 TNLs, 555 CNLs, and 163 RNLs. It is obvious that the four types of NLRs are unevenly distributed among different accessions (Figure 1, Supplemental Table S2). Based on the mean values, the numbers of RNLs are the most stable with roughly 10 possessed by each accession. This indicates that the copy numbers of RNLs are the most conservative. The numbers of CNLs in each accession are more marked different, with only 17 genes in PCA accession (Pak choi) genome, and 51 genes in TBA accession from Tibet, China. The large copy number of CNLs in TBA accession might be adaptive to the special environment of Tibet. Similarly, the numbers of TNLs are significantly different in different accessions. Most of accessions encode only dozens of TNLs, such as CCA accession (Chinese cabbage), TBA accession (From Tibet, China), while some accessions encode over 100 TNLs, such as Chiifu (Chinese cabbage), OIA (Oil seeds) and MIZ (Mizuna). This suggests that TNLs are specifically expanded in certain accessions. Overall, the numbers of different NLR types vary dramatically, and different *Brassica rapa* accessions contain different combination pattern of four types of NLRs, which indicates the functional diversity of NLRs.

**Figure 1.**
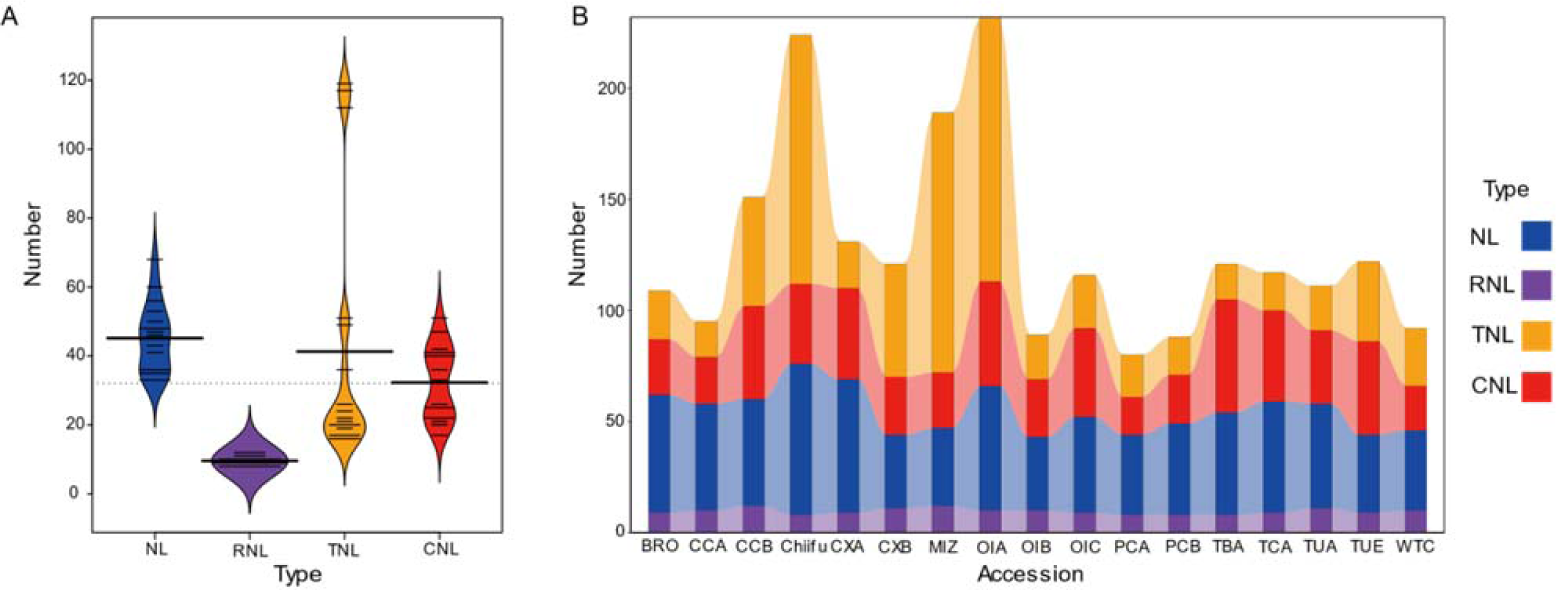
Numbers of four types of NLRs in 17 *Brassica rapa* accessions. A) The distribution of four types of NLRs in 17 *Brassica rapa* accessions. Four thickened solid black lines represent the median of four types of NLRs. Multiple thin solid black lines represent the original data points. Single long dotted line indicates the mean value of all data. Blue, purple, orange, red colors represent NLs, RNLs, TNLs, and CNLs, respectively. B) Stacked bar plot of four types of NLRs in 17 *Brassica rapa* accessions. NLs, RNLs, TNLs, and CNLs are colored blue, purple, orange, and red, respectively.

NLRs identified from 13 *Brassica rapa* accessions with chromosome information in this study are mapped to the chromosomes of each *Brassica rapa* accession according to their physical positions (Figure 2). Interestingly, besides the stable copy numbers, RNLs physical positions are more conserved than the other three types of NLRs, especially in the chromosome A01, A02, A06, A07, A08, A09, and A10. The stable copy number and physical positions among the different *Brassica rapa* accessions demonstrate that RNLs might play a relative conserved and important role in *Brassica rapa* pathogen resistance. On the contrary, the physical positions of TNLs and CNLs are varied in different accessions. For example, at the beginning of A02 and A03 chromosomes, TNLs form clusters in Chiifu (Chinese cabbage), and MIZ (Mizuna), but other accessions do not contain this kind of TNL clusters at the same position, which shows the specific expansion in certain accessions. The specific expansion of TNLs and CNLs indicates their specific biofunctions in the related subspecies.

**Figure 2.**
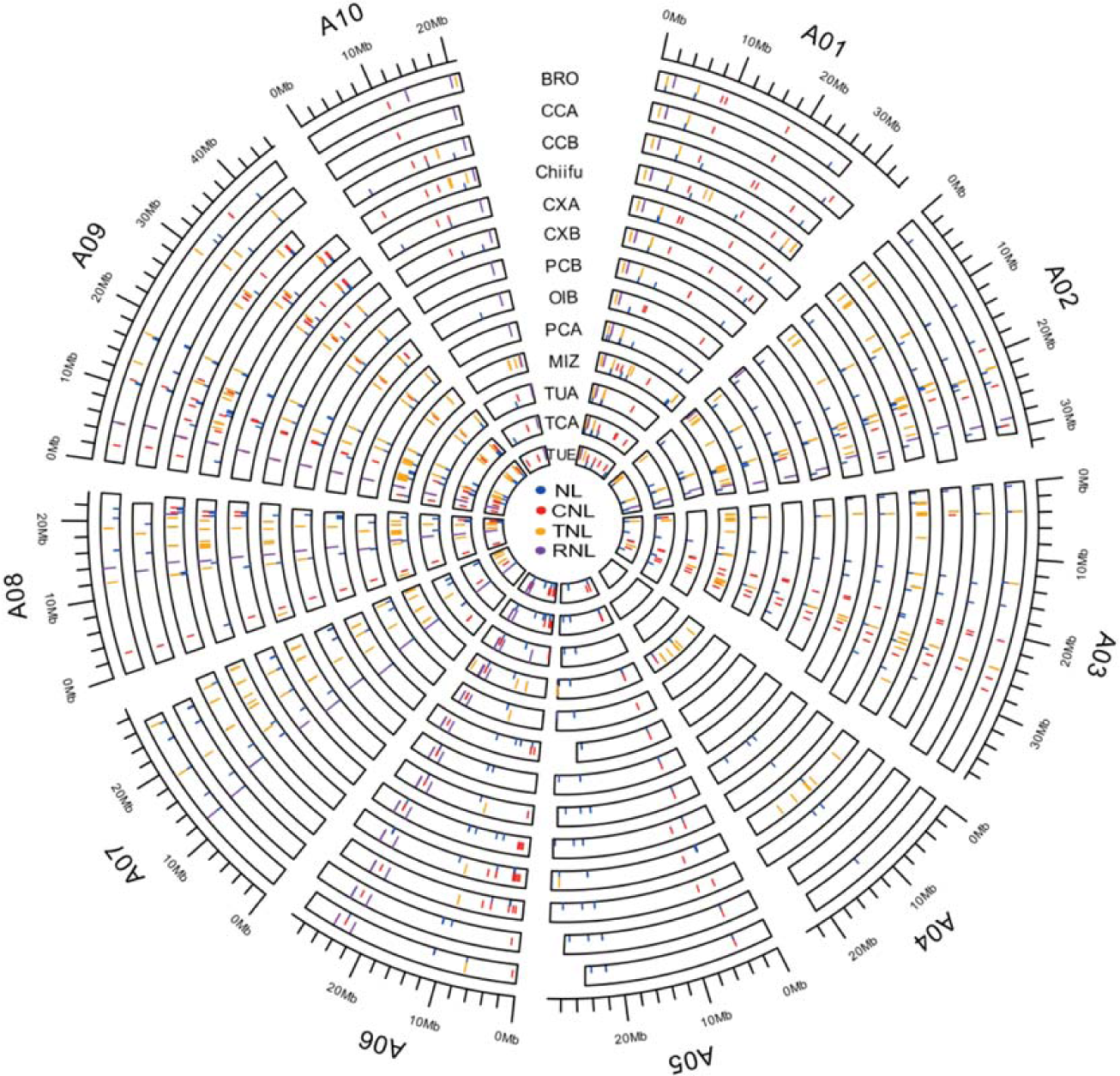
The physical locations of NLRs from 13 *Brassica rapa* genomes. NLs, RNLs, TNLs, and CNLs are colored blue, purple, orange, and red, respectively.

Though the positions of RNLs are relatively stable, there are 3 RNLs (*A06p15500.1_BraTUA*, *A08p29820.1_BraCCB*, and *A10p18270.1_BraBRO*) that only exist in one accession (Figure 2). *A06p15500.1_BraTUA* located on chromosome A06 of TUA (Turnip) belongs to OG0000034 ortholog group mainly containing the RNLs located on A10, except three RNLs without the chromosome information. Transposable elements (TEs) might be the reason for *A06p15500.1_BraTUA* translocation*. A08p29820.1_BraCCB* located on chromosome A08 of CCB (Chinese cabbage) is singleton and the most parsimonious explanation for this is *de novo* origination.

### The Pan-NLRomes of *Brassica rapa*

The pan-genome of a certain species represents the entire genetic information of that species, then pan-NLRomes are the NLR components of a pan-genome (Barragan and Weigel, 2021). As NLRs cannot be captured fully from a single genome, the pan-NLRomes can provide a complete inventory of all NLRs in a certain species. All the NLRs identified form this study are clustered into 175 ortholog groups (OGs) containing at least two genes based on the sequence similarity and phylogenetic relationships. Only a little more than 3% of all NLRs (7 NLRs) are singletons. 95% of all the 175 OGs could be found in about 9 randomly chosen accessions (Figure 3A). This closed pattern demonstrates that the NLRs discovered in this study is saturated. All the 175 NLR OGs were classified into three categories (Core, Shell, and Cloud) based on the distribution of their NLR members. 39 OGs (22.3%) were defined as core NLRomes, comprising 1,030 (47%) genes from at least 16 accessions. 63 OGs (36%) were defined as clould NLRomes, comprising 227 (10.4%) genes from 3 or fewer accessions. The last categories shell NLRomes were defined with 73 OGs (41.7%), containing 924 (42.2%) genes that were found in at least 4 but fewer than 16 accessions (Supplemental Table 3). Notably, 139 of all the 163 RNLs (85.8%) belong to the core categories, and 160 TNLs account for 70.5% in the cloud categories (Figure 3B, Supplemental Table 4). These two kinds of unbalanced distribution of RNLs and TNLs are consistent with their numbers and physical locations in different *Brassica rapa* accessions. In general, core NLRs perform basic biological functions, especially in the aspects of growth and development of plants. The other NLRs from the shell and cloud categories might be the results of accession-specific loss or expansion during evolution. They are the main force in improving the diversity of Pan-NLRomes, and might play essential roles in fighting with certain pathogens.

**Figure 3.**
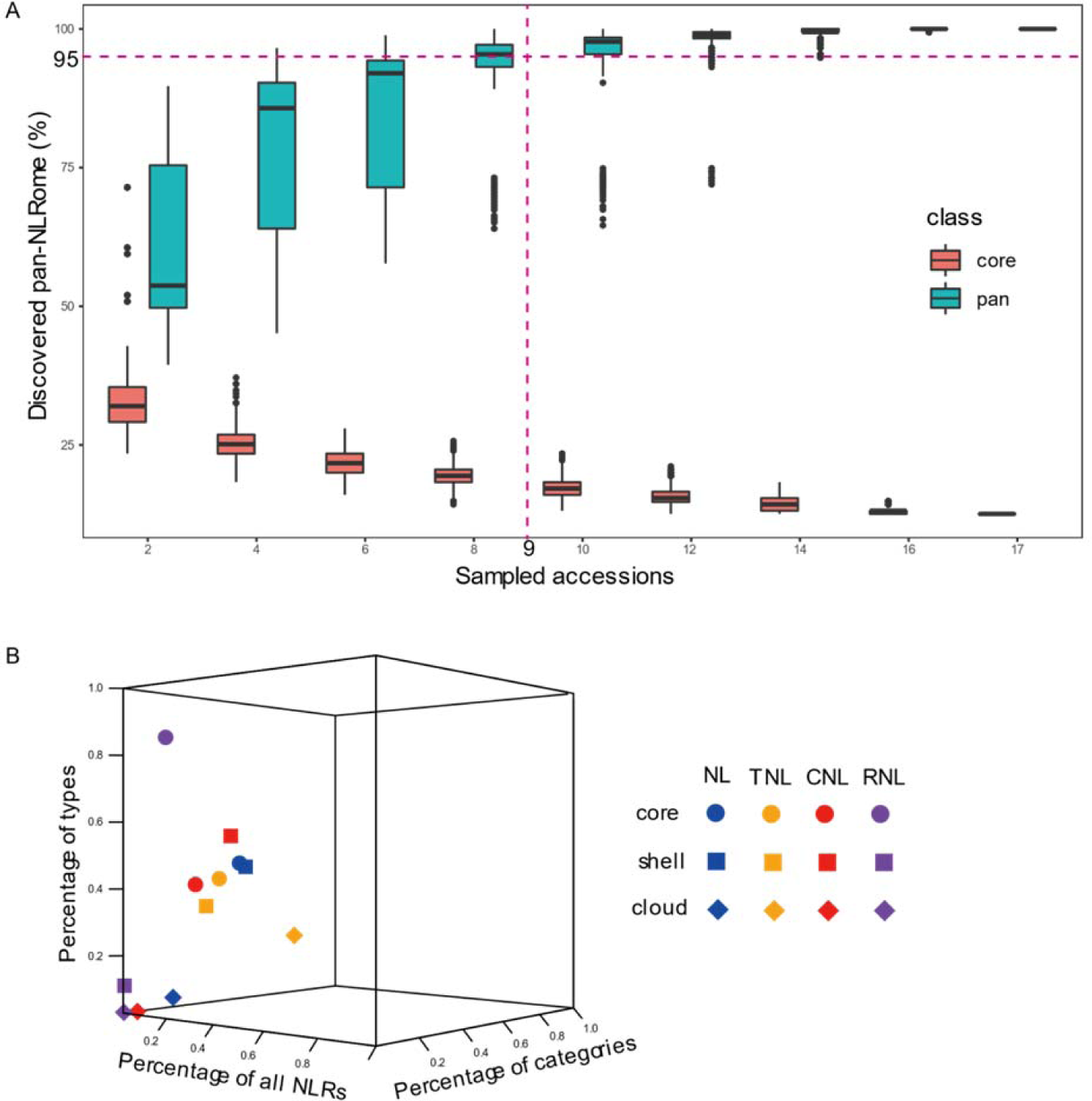
The pan-NLRome distributions and proportions. A) The saturation of pan-NLRome discovery of 17 *Brassica rapa* accessions. Blue boxes indicate the percentage of pan-NLRome recovered from 17 randomly selected accessions of different sizes (with 1,000 bootstraps). Red boxes indicate the core-NLRome present in all selected accessions (with 1,000 bootstraps). Horizontal dashed line indicates 95% of pan-NLRome discovered. Vertical dashed line indicates that 95% of the pan-NLRome can be recovered with 9 accessions (1,000 bootstraps). B) Percentages of the different type-category NLRs in all NLRs, different types, and different categories, respectively. NLs, RNLs, TNLs, and CNLs are colored blue, purple, orange, and red, respectively. Circles represent core NLRs, squares represent shell NLRs, and rhombuses represent cloud NLRs. For example, blue circles represent the NLs belonging to core category.

### Integrated domain diversity of NLRs

Besides the three canonical domains, integrated domains (IDs) can dramatically increase the diversity of NLRs. 365 NLRs contain at least one IDs, accounting for up to 16.7 percent of all NLRs of this study (Supplemental Table 3). There are 93 distinct Pfam domains contained by the 365 NLRs (Supplemental Table 3). All the 93 IDs were compared in the three categories (Figure 4A) and the four different types (Figure 4B). There is only one domain (ATPase family associated with various cellular activities, PF00004) contained by all the three categories. Notably, NLRs of shell category carry 35 category-specific IDs, which is largest amount compared with that of the other two categories. In the aspect of NLR different types, none of ID is shared by all the 4 types of NLRs. All the RNLs only have 4 different IDs distributed in 6 accessions. Moreover, RNLs have no type-specific IDs. On the contrary, NLs, TNLs, and CNLs have 30, 20, and 15 type-specific IDs, respectively. Taken altogether, distinct IDs can improve the diversity of NLRs, especially in the pathogen effector recognition.

**Figure 4.**
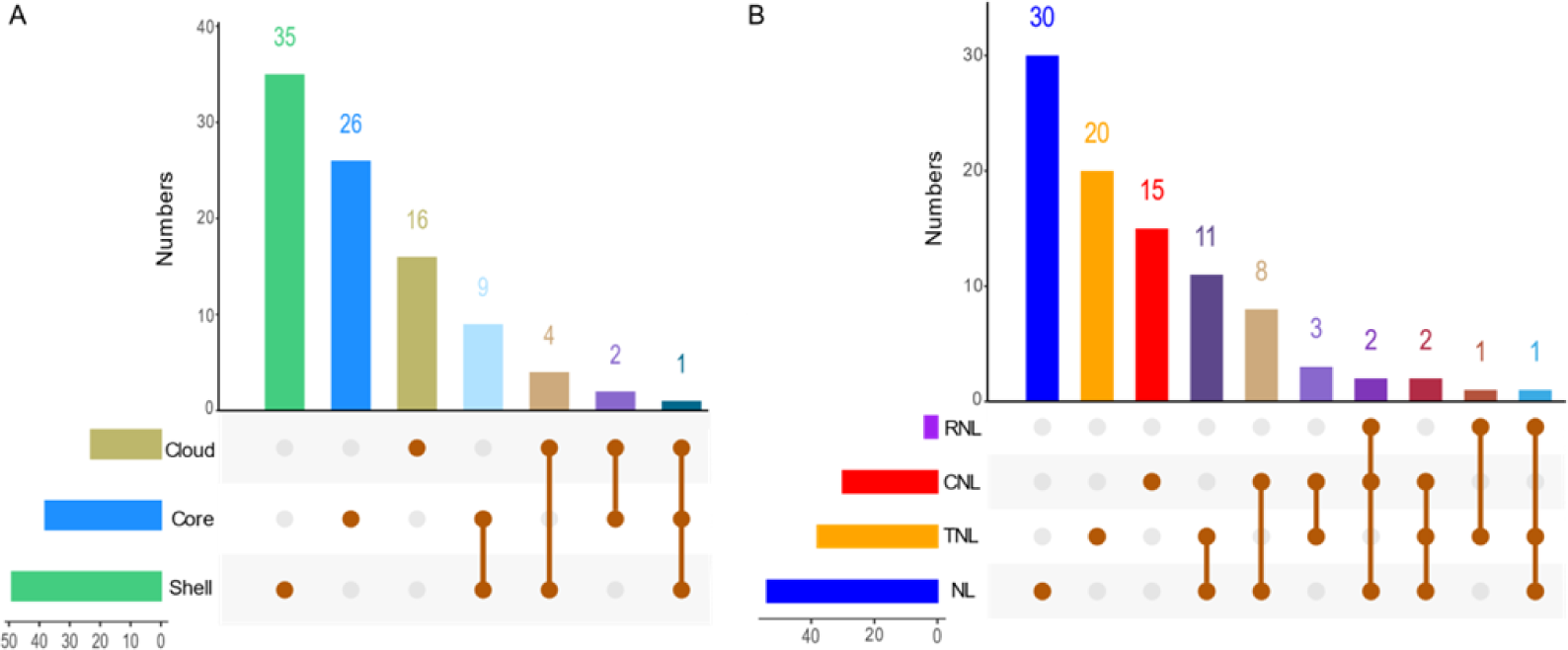
Upset intersection of IDs of different categories (core, cloud, and shell) and different types (NLs, RNLs, TNLs, and CNLs). A) Upset intersection of IDs of different categories (core, cloud, and shell). B) Upset intersection of IDs of different types (NLs, RNLs, TNLs, and CNLs).

The top ten most commonly found IDs in *Brassica rapa* NLR proteins are AAA_14 (PF13173), CDC48 N-terminal domain (PF02359), VQ motif domain (PF05678), Calmodulin_bind (PF07887), zf-CCCH (PF00642), Rx N-terminal domain (PF18052), neprosin (PF03080), AAA_assoc (PF14363), B3 DNA binding domain (PF02362), and Omp85 superfamily domain (PF01103). Rx N-terminal domain and Calmodulin_bind domain are associated with the induction of plant defence responses. The CDC48 N-terminal domain is a substrate recognition domain that binds peptides, prevents protein aggregation, and catalyzes the refolding of allowed substrates (Bodnar and Rapoport, 2017). The Zinc finger domain can interact with the 3’ untranslated region of mRNA and has a deacetylating and degrading effect (Hall, 2005). In general, the biofunctions of IDs coordinate with disease-resistant function, such as providing energy, signal transduction, recognition of specific DNA sequences, etc. There are also IDs with unknown functions that may be associated with stability of protein structure or catalysis or inhibition, and further studies are needed to decipher their functions.

It is noteworthy that 75 NLRs contain RPW8 domain and CC domain at the same time, and 7 NLRs hold TIR domain and CC domain at the same time. All the RNLs contain varying degrees of random coil. It may be that some RNLs produced the random coil through nonsynonymous mutations or other kinds of molecular mechanisms during the evolution, then evolved into the initial type of CNLs. This hypothesis may explain the relatively unequal distribution of CNLs in *Brassica rapa*.

### Accession-specific expansion of TNLs

Among the four types of NLRs, TNLs have the largest number variation in each accession, and the difference in number can reach more than ten times. The distribution of TNLs on chromosomes is unequal compared to the other three types of NLRs (Figure 2). Since the NB-ARC domain (PF00931) is the most conserved domain, we extracted the NB-ARC domain of each NLRs for constructing a phylogenetic tree (Figure 5A). Based on the phylogenetic relationship, all the NLRs can be clustered into three monophyletic clades (Clade I, Clade II, and Clade III). Most RNLs mainly locate in the base of the phylogenetic tree with the relatively short branches. A part of NLs are also quite conserved, and they clustered in Clade I with most RNLs. On the contrary, CNLs and TNLs are more varied. Most CNLs and NLs locate in Clade II, and they do not cluster respectively. Clade III mainly contains TNLs, in which, the expansion of TNLs is particularly obvious in the subclade I which contains 286 members. Moreover, the branches of subclade I are relatively shorter than the branches of others in Clade III, which implies that the expansion happened in a short period of time.

**Figure 5.**
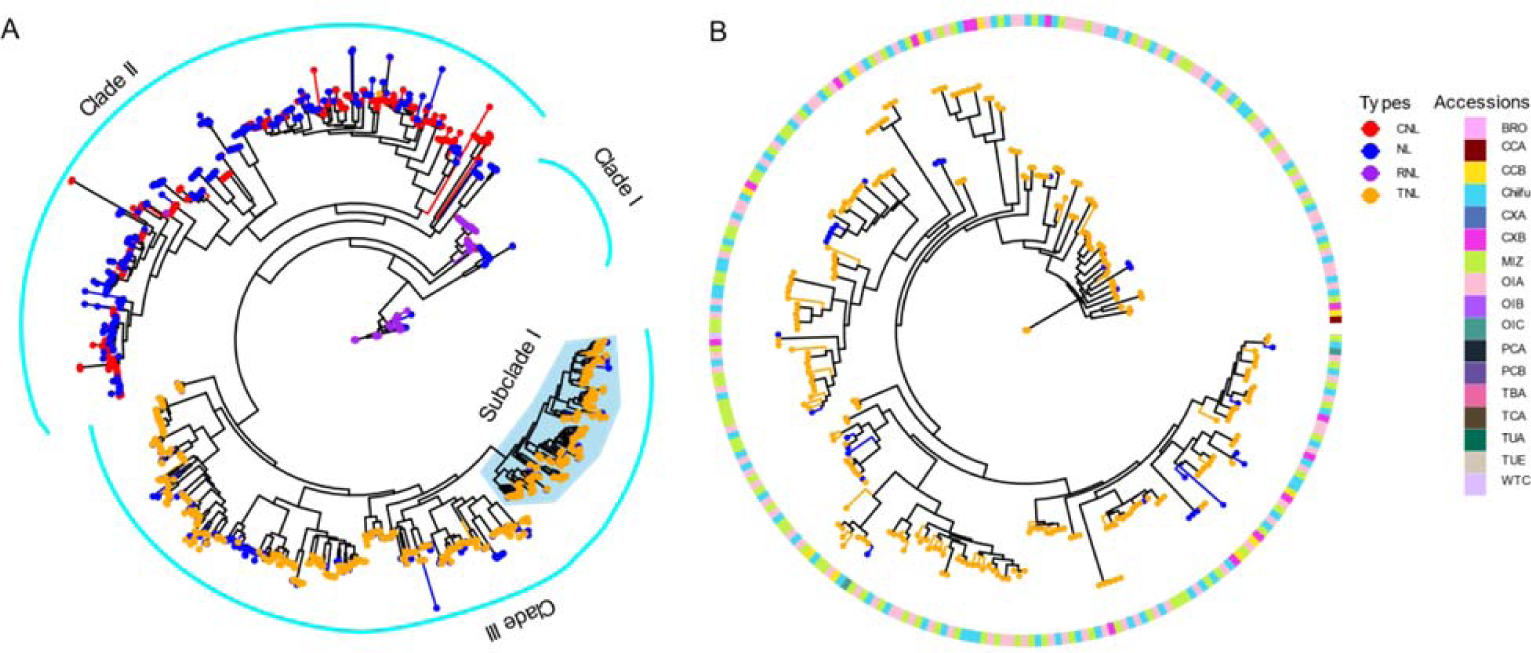
Maximum-likelihood phylogeny of NLRs based on NBS domain. A) Maximum-likelihood phylogeny of all the NLRs with different tip colors. NLs, RNLs, TNLs, and CNLs are colored blue, purple, orange, and red, respectively. B) Maximum-likelihood phylogeny of subclade I from Clade III in Figure 5A, and the outermost circle represents the *Brassica rapa* accessions information of each tip.

In order to deeply insight into the evolution of subclade I of Clade III, NLRs were extracted from subclade I, and phylogenetic tree was built based on the NB-ARC domain (Figure 5B). In consideration of the accession information, the numbers of TNLs from OIA (Oil seeds), MIZ (Mizuna), and Chiifu (Chinese cabbage) are the top three in subclade I of Clade III, with 90, 86, and 82, respectively (Supplemental Table S5). That is to say, NLRs from those three accessions account for up to ninety percent of subclade I, which provides strong evidence for TNL accession-specific expansion. In the aspect of three different NLR categories, subclade I contains one gene that is singleton. The remaining 285 genes contain 173 genes belonging to the cloud category and 112 genes belonging to the shell category. Most notably, there is no one belonging to the core category (Supplemental Table S5). Another thing to be aware of is that the majority of NLR proteins of subclade I do not contain IDs and some of them contain AAA domain which might suggest that NLRs in this clade were newly originated. All the clues mentioned above signify that the Clade III NLRs, especially the TNLs from subclade I are newly expanded in some certain accessions when facing the challenges of pathogens or other biotic stresses. It is worth studying their biofunctions and illuminating their molecular mechanism in the future.

### Positive selection sites of NLRs

RNLs are relatively conserved in the aspects of location and numbers in different *Brassica rapa* accessions. On the contrary, TNLs, especially in the subclade I of Clade III experienced accession-specific expansion. We performed positive selection analyses to tested whether RNLs and accession-specific expanded TNLs underwent adaptive evolution. RNLs mainly group into 6 OGs. Positive selection analyses showed that the numbers of their positive selection sites range from 1 to 21 (Figure 6, Supplemental Table S6). OG0000006 and OG0000034 only have 3 and 1 positive selection sites, respectively. They are all located outside of the conserved structural domains. The other four OGs have 16, 21, 17 and 14 positive selection sites, respectively. Unlike the first two OGs, their positive selection sites are mainly located in the RPW8 domain. The OGs from subclade I of Clade III have more positive selection sites than that of RNLs, with an average of 38 selection sites of each OG (Figure 6, Supplemental Table S6). Moreover, their positive selection sites are mainly distributed in intrinsically disordered regions and IDs. All in all, RNLs and accession-specific expansion TNLs all underwent adaptive evolution, but the positive selection sites which they contain are dramatically different. As the positive selection sites play important roles in recognition of pathogen effectors, the accession-specific expansion TNLs might be on the front line in fighting against the diverse and fast-evolving pathogens, which represents an intriguing evolutionary strategy in the plant-pathogen arms race, while RNLs might not directly interact with pathogens.

**Figure 6.**
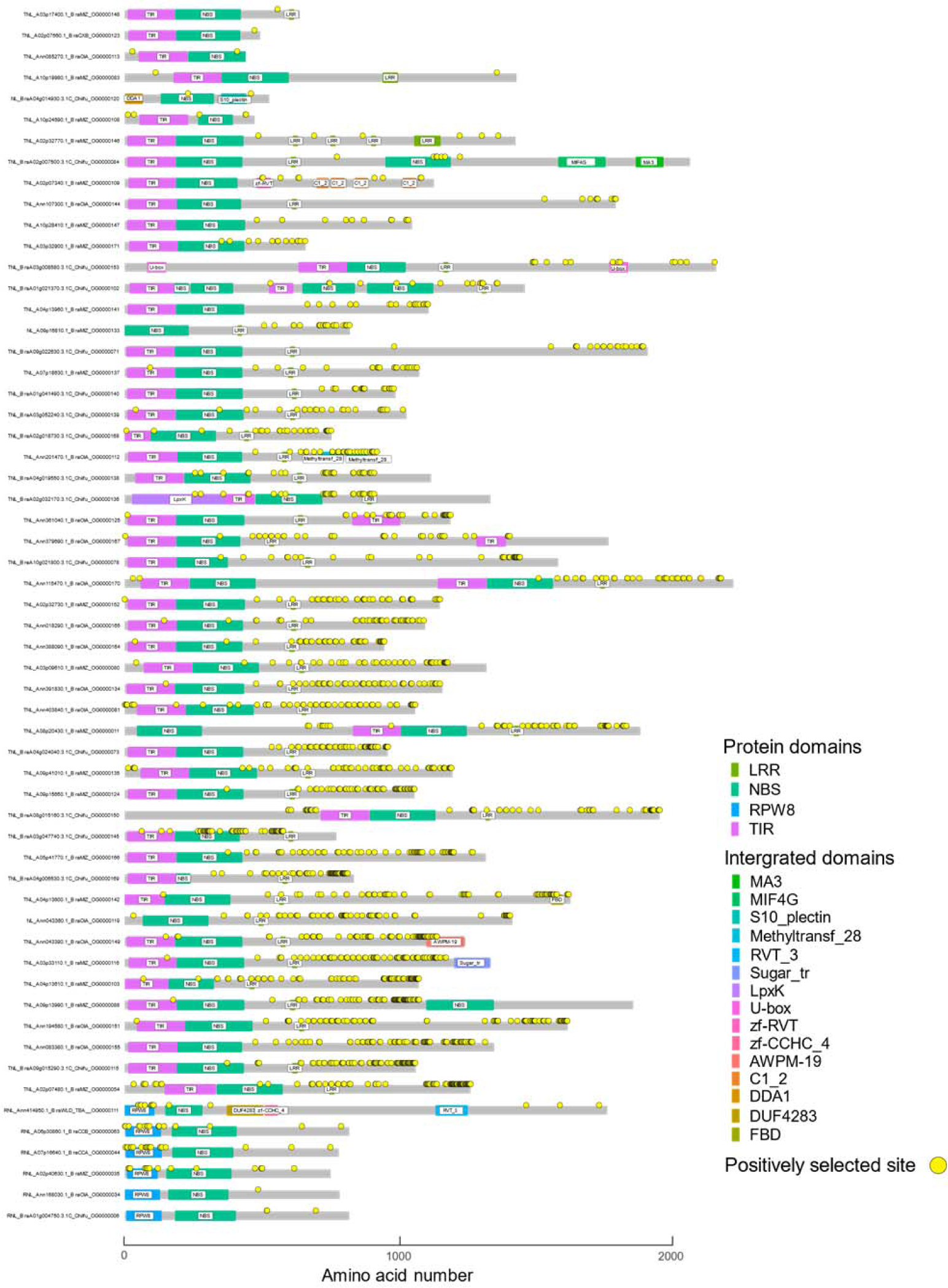
Positive selection sites and protein schematics of representative TNLs and NLs from the accession-specific expansion TNL-type OGs and NL-type OGs, and representative RNLs of six RNL-type OGs. The detailed information for each gene and OG is available in Supplemental Table S5.

### Protein structure analyses and binding capacity prediction

In order to illustrate the location of positive selection sites in 3D protein structures, 3 representative RNL-type proteins (BraA01g004750.3.1C from OG0000006, A02p40630.1_BraMIZ from OG0000035, and Ann414950.1_BraWLD from OG0000111) and 3 accession-specific expanded TNL-type proteins (A08p20430.1_BraMIZ from OG0000011, BraA09g022630.3.1C from OG0000071, and Ann391830.1_BraOIA from OG0000134) were selected to compute their 3D structure by Alphafold2 (Figure 7, Supplemental Figure S1). Most of the positive selection sites are located on the surface of 3D protein structures (Figure 7, A and D, Supplemental Figure S1, A, C, E, and G), where they may play roles in interaction with pathogen effectors. It’s noteworthy that the 3D structure of TNL-type protein (Ann391830.1_BraOIA, Figure 7D) contains a “ring”, and the positive selection sites of this protein are mainly distributed in the inner side surface of the “ring”, which are the strong evidence for pathogen effector recognition. We also predicted the sites binding probabilities of the 3 RNLs and 3 TNLs by a deep learning network (Figure 7, B and E, Supplemental Figure S1, B, D, F, and H). The binding probabilities of positive selection sites were extracted, and the binding probabilities higher than 0.5 were highlighted (Supplemental Table S7). There are 1 RNL-type protein (Ann414950.1_BraWLD) and 2 TNL-type proteins (A08p20430.1_BraMIZ and BraA09g022630.3.1C) possessing more than half percent of positive selection sites with the binding probabilities higher than 0.5. On the contrary, most of the positive selection sites distributed in the inner side surface of the ring from Ann391830.1_BraOIA have lower binding probabilities (Supplemental Table S7). This counterintuitive result should be tested through site binding capacity experiments in the future. On the whole, the combination of positive selection sites, binding probabilities, and 3D structure results provide some clues about the important amino acid sites of NLRs for pathogen recognition. The amino acid sites with obvious positive selection signals, high binding probabilities, and distribution in certain regions, such as ‘package’ and ‘ring’ might be the candidate sites for functional innovation in facing with the rapidly evolved pathogen.

**Figure 7.**
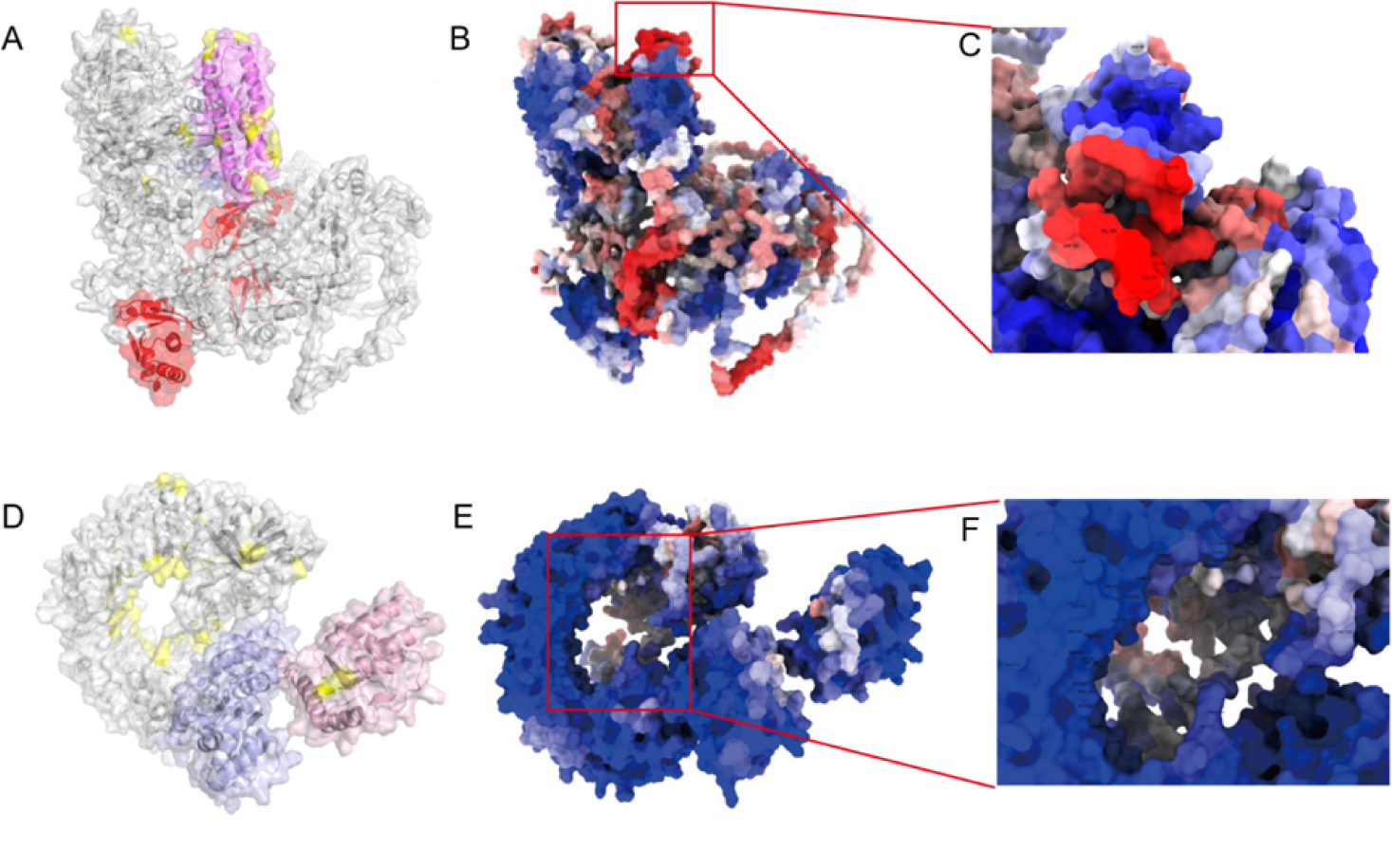
3D structures of representative RNL and TNL proteins, and their binding capacity prediction. A) The 3D structure of representative RNL protein (Ann414950.1_BraWLD_OG0000111). NBS domain, RPW8 domain, and integrated domain regions are shown in light blue, violet, and red, respectively. Positive selection sites are colored yellow. B) Protein binding sites of Ann414950.1_BraWLD_OG0000111 predicted by deep learning network. Red denotes a high binding probability, and the blue indicates a low binding probability. C) Enlarged view of the high binding probability region with positive selection sites from Figure 7B. Red denotes a high binding probability, and the blue indicates a low binding probability. D) The 3D structure of representative accession-specific expanded TNL protein (Ann391830.1_BraOIA_OG0000134). NBS domain and TIR domain are shown in light blue and light pink, respectively. Positive selection sites are colored yellow. E) Protein binding sites of Ann391830.1_BraOIA_OG0000134 predicted by deep learning network. Red denotes a high binding probability, and the blue indicates a low binding probability. F) Enlarged view of the “ring” region with positive selection sites from Figure 7E. Red denotes a high binding probability, and the blue indicates a low binding probability.

## Discussion

Previous study showed that *NLR* gene numbers vary among *Brassica napus* (641), *Brassica rapa* (249), and *Brassica oleracea* (443) (Alamery et al., 2018). In another study, 204 *NLR* genes were identified in *Brassica rapa* through BLAST and HMMsearch methods (Zhang et al., 2016). The numbers of *NLR* genes identified in the 17 *Brassica rapa* accessions range from 80 to 232, which is consist with the previous studies. There are at least two reasons that influence the accuracy of the *NLR* gene identification. The first reason is the mining method. The second is the quality of the genome assembly and genome annotation. As some *NLR* genes are clustered together, it brings on challenges to assembly this region. Additionally, gene model prediction cannot find all the *NLR* genes for the reason that some resistant genes only express when facing pathogens. *De novo* mining *NLR* genes from the whole genome sequence based on the deeply studied *NLR* gene sequences through machine learning might be the promising solution.

NLRs harbor additional, non-classical IDs (Kourelis and van der Hoorn, 2018). The different types of IDs can increase the diversity of NLRs. Those IDs may play a role in oligomerization, downstream signaling or perform other functions (Read et al., 2020). In the head-to-head paired *NLR* genes, one acts as an executor and the other acts as a sensor. The sensor often has an ID which might be a bait for the effectors of pathogens (Barragan and Weigel, 2021). 93 IDs were identified in the pan-NLRomes of *Brassica rapa*, the functions of which are mainly associated with DNA binding such as B3 binding domain and protein-protein interaction such as zinc finger C-x8-C-x5-C-x3-H type domain, suggesting that certain protein domains combined with canonical NLR protein domains may be evolutionarily advantageous. As minor nonsynonymous mutations in the ID region can expand the response profile to some pathogens (Barragan and Weigel, 2021), all the IDs mined from the pan-NLRomes of *Brassica rapa* can act as the candidates for genetic manipulation through gene editing and so on.

The numbers of RNLs are most stable and the phylogenetic tree shows a conserved evolutionary pattern, which means RNLs play the most fundamental role in *Brassica rapa* over a long period of evolution and differentiate a more comprehensive disease resistance gene system. Phylogenetic analysis of *NLR* genes in five Brassicaceae genomes showed that RNLs is the an ancient *NLR* genes with conserved function (Zhang et al., 2016). RNLs have not experienced dramatically expansion like TNLs in *Brassica rapa* pan genomes, which is consist with the *Arabidopsis thaliana*, *Carica papaya*, and *Thellungiella salsuginea* (Zhang et al., 2016; Van de Weyer et al., 2019). As those species are all from Brassicales, further studies should be conducted to test whether the conserved evolution pattern of RNLs is prevalent out of Brassicales, especially on the pan-genome level. It is likely that RNLs are not in the manner of gene-for-gene in pathogen recognition, but they assist other *NLR* genes to accomplish a resistance response (Xiao et al., 2001). Therefore, it is not necessary for RNLs to expand their numbers for pathogen recognition in plants (Tirnaz et al., 2020). The main difference between the four types of *NLR* genes (TNL, RNL, CNL, and NL) is the N-terminal domain. An interesting and important question comes out that does the RPW8 domain give the distinct function to RNLs? More experiments should be conducted to elucidate the uncertainty in the future.

Contrary to the conserved evolutionary pattern of RNLs, *TNL* genes specifically expanded mainly in three *Brassica rapa* accessions and they accumulated more positive selection sites. The TNL numbers of the 17 *Brassica rapa* accessions vary between 16 to 119 (Supplemental Table 2), which may hint that TNLs are in the manner of gene-for-gene in pathogen recognition. Their large number of positive selection sites might be the result of adaptation to the rapid pathogen evolution. The specifically expanded TNLs might play significant roles in facing with certain pathogens in the three *Brassica rapa* accessions. Previous studies show that TNLs might participate in resisting clubroot disease (Hatakeyama et al., 2013; Hatakeyama et al., 2017). Clubroot is one of the most serious diseases of many important cruciferous vegetables and oilseed crops worldwide, and more than 1/3 of the area planted to cruciferous crops is infested with clubroot year-round (Yang et al., 2022). Hatakeyama et al. (2013) showed that the Crr1a protein encoded by TNL type *NLR* gene inhibits the development of plasmodia during the secondary infection phase. They further clarified the main clubroot resistant locus *CRb* in Chiifu comprising at least six ORFs similar to TNL type *NLR* genes, which are in tandem with the same orientation (Hatakeyama et al., 2017). But there are no reports about the functions of the expanded TNLs in mizuna (MIZ) or oil seeds (OIA). We will next investigate the biofunction of the expanded TNLs in these three accessions under the challenges of clubroot pathogens and other possible pathogens. Though the specific expansion of TNLs exist in some dicots, they are absent from all the monocots. On the contrary, the other three types of *NLR* genes are present in almost all the sequenced Angiosperms (Yue et al., 2012). Taken together, the evolutionary mechanisms of different types of *NLR* gens are very complex.

## Materials and Methods

### Plant materials and data collection

The protein sequences and annotation files of 17 *Brassica rapa* accessions were retrieved from previously published works (Cai et al., 2021, Supplemental Table 1). The CDS sequence files of *Arabidopsis thaliana*, *Capsella rubella*, *Schrenkiella parvula, Sisymbrium irio, Brassica oleracea*, *Brassica napus,* and *Brassica juncea* were downloaded from Brassicaceae Database (http://brassicadb.cn, Chen et al., 2022).

### NLRs Identification and protein architectures prediction

The Hidden Markov Model profiles of NB-ARC (PF00931), TIR (PF01582), and RPW8 (PF05659) domains were retrieved from the Pfam database (http://pfam-legacy.xfam.org/, Mistry et al., 2020). HMMER v3.2.1 (Eddy, 2011) were employed to search against the whole genomes of 17 *Brassica rapa* accessions with a cutoff E-value 10^−3^. Coiled-coil (CC) domains were predicted by deepcoil python package utilizing the Keras machine learning library (Ludwiczak et al., 2019). Finally, all the obtained NLR candidates were classified into the NL, TNL, CNL, and RNL types by custom python scripts. The integrated domains of all the NLR proteins were predicted by Interproscan software (Jones et al., 2014) which integrates multiple conserved protein domain databases.

### Sequence alignment and phylogenetic analyses

NB-ARC domains of all the NLR proteins identified from this study were extract by custom python scripts. Then they were aligned by Mafft with default parameters (‘--auto’) (Katoh and Standley, 2013). The best-fitting model of the multiple sequence alignments of NB-ARC domains was determined by ModelFinder (-m MFP, Kalyaanamoorthy et al., 2017). The maximum likelihood (ML) phylogenetic trees were constructed using IQ-TREE (Nguyen et al., 2015) with 1,000 bootstrap replicates. Ggtree R package was used to plot and annotate the phylogenetics trees (Yu, 2020).

### Orthogroups (OGs) identification

OrthoFinder (Emms and Kelly, 2015) was used to identify Orthogroups (OGs) in 17 *B. rapa* accessions. The OGs were grouped into three categories (core, shell, cloud) according to previously published works (Van de Weyer et al., 2019). We divided the OGs into NL, CNL, TNL and RNL types if they account for more than or equal to 60% of each OG.

### Positive selection analyses and of *NLR* genes

The proteines from 6 RNL-type OGs and 55 accession-specific expansion TNL-type OGs were used as queries to search against seven other species (*Brassica oleracea*, *Brassica napus*, *Brassica juncea*, *Arabidopsis thaliana*, *Schrenkiella parvula*, *Capsella rubella*, *Sisymbrium irio*, Supplemental Table 1) by blastp algorithm (Altschul et al., 1990). The coding sequence (CDS) of the *NLR* genes were aligned using MEGAX (Kumar et al., 2018). Phylogenetic trees were constructed by selecting the CDS of the best aligned genes in each species. The amino acid positive selection sites were calculated using codeml in the PAML package (Xu and Yang, 2013). M7 (beta) and M8 (beta&) models were compared to detected the evidence of positive selection based on the likelihood ratio test through the Bayes empirical Bayes (BEB) procedure. Positive selection sites and protein schematics were visualized by drawProteins R package (Brennan, 2018)

### 3D structure computation and binding probabilities prediction

The 3D structures of NLR proteins were generated by AlphaFold2 (Jumper et al., 2021), and the best model determined by the pLDDT score was selected. The 3D structures were visualized by PyMOL (https://pymol.org/2/). The protein binding probabilities of NLR proteins were predicted by ScanNet web server (http://bioinfo3d.cs.tau.ac.il/ScanNet/index_real.html, Tubiana et al., 2022) and the outcomes were depicted by ChimeraX 1.6.1 (Pettersen et al., 2021).

## Acknowledgments

This work was supported by the Scientifc Research Project of Introducing Talents of Hebei Agricultural University (No. YJ201944); the Innovative Research Group Project of Natural Science Foundation of Hebei Province (Grant No. C2020204111).

## Author contributions

C.W. and X.S. designed the research. C.W. improved the manuscript. S.L. and Y.Z. performed research, analyzed data and wrote the manuscript. B.K. and J.W. performed the data analyses. X.P., S.Z., R.Z., H.W., J.L., and J.L. analyzed partial data; Y.Z., T.H., S.W., Z.L., L.W. collected data.

## Supplemental materials

**Supplemental Figure S1.** 3D structures of representative RNL and TNL proteins, and their binding capacity prediction. NBS domain, RPW8 domain, and TIR domain regions are shown in light blue, violet, and light pink, respectively. Positive selection sites are colored yellow. Red denotes a high binding probability, and the blue indicates a low binding probability. A) and B) 3D structure of representative RNL protein (Chiifu_BraA01g004750.3.1C_OG0000006) and its binding capacity prediction. C) and D) 3D structure of representative RNL protein (A02p40630.1_BraMIZ_OG0000035) and its binding capacity prediction. E) and F) 3D structure of representative accession-specific expanded TNL protein (Chiifu_BraA09g022630.3.1C_OG0000071) and its binding capacity prediction. G) and H) 3D structure of representative accession-specific expanded TNL protein (A08p20430.1_BraMIZ_OG0000011) and its binding capacity prediction.

**Supplemental Table S1**. Lists of 17 *Brassica rapa* accessions (Cai et al., 2021), and 7 other species of Brassicaceae used in this study.

**Supplemental Table S2**. The total number and four types of NLRs identified in 17 *Brassica rapa* accessions.

**Supplemental Table S3**. Lists of all the NLRs identified in 17 *Brassica rapa* accessions with gene name, accession, type (NL, TNL, RNL, and CNL), category (core, cloud, and shell), and integrated domain information.

**Supplemental Table S4**. Percentages of the different type-category NLRs in all NLRs, the same categories and the same types. Two dramatically unbalanced proportions (TNL-cloud and RNL-core) are highlighted.

**Supplemental Table S5**. The list of NLRs in subclade I from clade III with gene name, physical position, integrated domain, and category information.

**Supplemental Table S6**. Positive selection sites of representative RNLs from six OGs and 55 representative accession-specific expanded TNLs and NLs from subclade I of Clade III.

**Supplemental Table S7**. Binding site probabilities of the positive selection sites from 3 representative RNLs and 3 representative accession-specific expanded TNLs. The binding probabilities higher than 0.5 are highlighted.

